# Histone lysine crotonylation regulates cell-fate determination of neural stem/progenitor cells by activating bivalent promoters

**DOI:** 10.1101/2020.05.27.118596

**Authors:** Shang-Kun Dai, Pei-Pei Liu, Hong-Zhen Du, Xiao Liu, Ya-Jie Xu, Cong Liu, Ying-Ying Wang, Zhao-Qian Teng, Chang-Mei Liu

**Affiliations:** State Key Laboratory of Stem Cell and Reproductive Biology, Institute of Zoology, Chinese Academy of Sciences, Beijing 100101, China; Savaid Medical School, University of Chinese Academy of Sciences, Beijing 100049, China; Institute for Stem Cell and Regeneration, Chinese Academy of Sciences, Beijing 100101, China

## Abstract

Histone lysine crotonylation (Kcr), an evolutionarily conserved and widely expressed non-acetyl short-chain lysine acylation, plays important roles in transcriptional regulation and disease development. However, its genome-wide distribution, correlation with gene expression, and dynamic changes during developmental processes are largely unknown. In this study, we find that histone Kcr is mainly distributed in active promoters, has a ge-nome-wide positive correlation with transcriptional activity, and regulates transcription of genes participating in metabolism and proliferation. Moreover, elevated histone Kcr activates bivalent promoters to stimulate gene expression in neural stem/progenitor cells (NSPCs), through increasing chromatin openness and recruitment of RNA polymerase II (Pol II). Functionally, these activated genes remodel transcriptome and promote neuronal differentiation.

**Author summary:** Overall, histone Kcr marks active promoters with high gene expression and modifies the local chromatin environment to allow gene activation, which influences neuronal cell fate. It may represent a unique active histone mark involved in neural developmental regulation.

## Introduction

Histone post-translational modifications (PTMs) occur on flexible amino-terminal tails protruding from nucleosome and globular histone cores. They form a scaffold around which DNA is wrapped, and regulate processes such as gene expression, DNA replication, DNA damage response, and higher-order chromatin structure [1]. Among these PTMs, histone lysine acetylation (Kac) at DNA regulatory elements is well-characterised and facilitates nucleosome eviction through disturbing interactions between histone and DNA. In addition, it promotes binding of Pol II to proximal promoters through recognition by effector proteins [2–4]. Recently, several other no-acetylation short-chain lysine acylations such as crotonylation (Kcr), propionylation, butyrylation, 2-hydroxyisobutyrylation, β-hydroxybutyrylation, succinylation, malonylation and glutarylation, have been found on both histones and non-histone proteins. Among them, histone Kcr has recently attracted increasing attention due to its crucial role in transcriptional regulation [5, 6].

Histone Kcr, specifically enriched at promoters and potential enhancers in the mammalian genome, has a stronger effect on transcription than histone Kac [7, 8]. Studies have implicated histone Kcr in development and diseases including self-renewal of mouse embryonic stem cells (mESCs), spermatogenesis, latency of human immunodeficiency virus (HIV), stress-induced depression, and protection from acute kidney injury [9–13]. Histone Kcr and Kac are believed to have the same transferases and “reader” modules. More concretely, acetyltransferases such as p300, CBP, and MOF catalyse Kcr on both histone and non-histone proteins [8, 14]. Among all known “reader” modules of histone Kac, YEATS domain of Taf14, YEATS2, and AF9 as well as DPF domain of MOZ and DPF2 display selective binding affinity for histone Kcr [15–18]. Class I histone deacetylases (HDAC I) and SIRT1 are major histone decrotonylases in mammals [10, 19, 20]. Furthermore, availability of intracellular crotonyl-CoA via crotonate treatment and inhibition of chromodomain Y-like (CDYL) can increase histone Kcr level [8, 11]. However, we know little about distribution and dynamic changes of histone Kcr at the genome level, which hinder cognition of roles of histone acylations in epigenetic and developmental studies.

As an active mark for transcription, histone Kac plays important roles in neurodevelopment and neurological diseases [21, 22]. Interestingly, a recent study showed that CDYL-mediated histone Kcr plays a critical role in regulating stress-induced depression [13]. Although histone Kcr is highly expressed and exists in combination with other histone PTMs in the mouse brain [11, 19, 23], its biological functions as well as regulatory mechanisms in neurogenesis are still unknown.

Here, we first investigated genome-wide distribution of histone Kcr and its correlation with gene expression. Next, we explored dynamic changes and regulatory mechanisms for transcription of histone Kcr in NSPCs. Finally, we performed integrated analysis with single-cell RNA sequencing data from embryonic cortex to illustrate biological functions of histone Kcr. Our results indicated that histone Kcr marks active promoters and can activate bivalent promoters by increasing chromatin openness, which stimulates gene expression and regulates NSPCs fate via remodeling transcriptome.

## Results

### Genome-wide distribution of histone Kcr and its correlation with gene expression

We first performed chromatin immunoprecipitation sequencing (ChIP-seq) using an antibody specifically recognising H3K9cr in E13.5 forebrain (S1A and B Fig), and found that H3K9cr was significantly enriched at active promoters marked by H3K4me3 and H3K27ac or H3K9ac (Fig 1A). K-means clustering divided annotated genes into distinct chromatin environments by H3K9cr signal (Fig 1B). Evidently, genes with higher H3K9cr level were usually accompanied with higher level of H3K4me3, H3K27ac and chromatin accessibility as well as lower level of H3K27me3 and DNA methylation (Fig 1B and 1C). Comparison of different combinations of histone PTMs revealed that H3K9cr marked genes with high levels of gene expression (Fig 1D). To further understand the general patterns of histone Kcr at the genome level, we analyzed publicly available ChIP-seq data studying histone Kcr in cancer cells [8, 19]. Our analysis showed unambiguously that H3K18cr, mainly located at active promoters was positively correlated with gene expression (S1C and S1D Fig). Importantly, despite considerable difference in histone Kcr level at promoters, there was a significant increase of transcription level with increasing histone Kcr signal (S1E Fig). Together, these results provided strong evidence that histone Kcr marks active promoters with high gene expression on a genome-wide scale.

**Fig 1.**
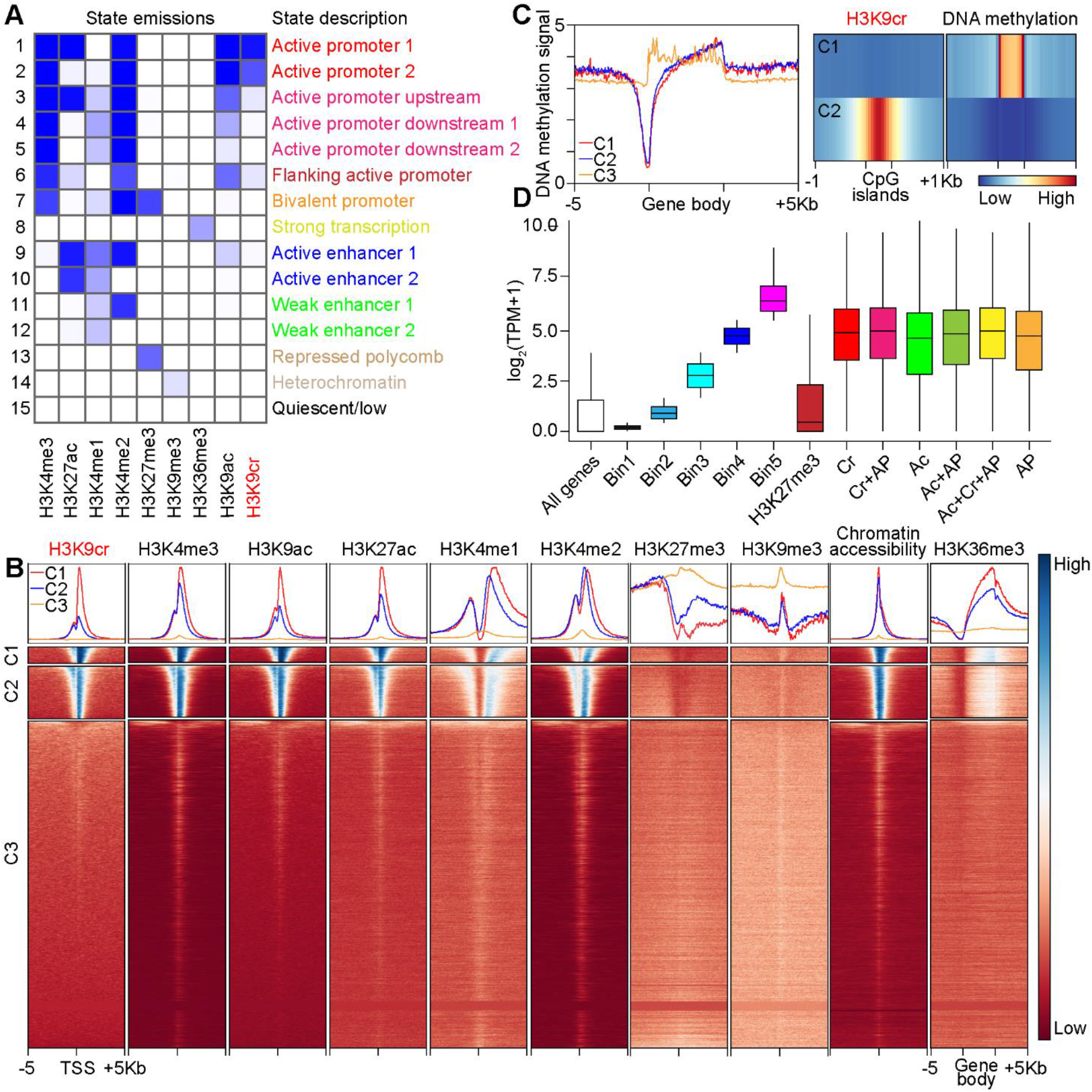
Genome-wide profiling and analysis of H3K9cr in the embryonic forebrain. (A) Heatmap showing distribution of H3K9cr signal on fifteen chromatin states modelled using ChromHMM, with each state description on the right panel. (B) Average profiles and heatmaps of histone marks, chromatin accessibility within +/-5 kb of transcription start sites (TSS) or gene body of genes which were divided to into three clusters and ranked from the highest to the lowest by H3K9cr signal (C1-C3). (C) Left panel, average profile of DNA methylation within +/-5 kb of gene body of three clusters of genes defined in B. Right panel, average heatmaps of H3K9cr and DNA methylation within CpG islands which were divided into two clusters by H3K9cr signal (C1-C2). (D) Boxplot showing gene expression changes of different groups of genes: Transcripts per million of genes were divided into five bins, as indicated, from lowly expressed genes (Bin1) to highly expressed genes (Bin5); genes under control of indicated histone marks (Cr: H3K9cr; Ac: H3K9ac; AP: H3K4me3+H3K27ac, active promoters).

### Functional annotation of histone Kcr peaks reveals its diverse functions

To investigate the biological functions of histone Kcr, we focused on H3K9cr peaks at promoter regions (Fig 2A and S2A Fig). These peaks were divided into two groups by H3K9cr signal (S2B Fig). De novo motif analysis of histone Kcr peaks discovered several transcription factors, such as TCF4 and SOX9, which are important regulators of neural development, thus implicating potential neural functions of histone Kcr (Fig 2B) [24, 25]. Further, we found that histone Kcr is primarily associated with genes involved in nucleic acid metabolism, protein quality control and cell proliferation (Fig 2C, S2C and S2D Fig), which highlights functional importance of histone Kcr in regulating basic cellular processes.

**Fig 2.**
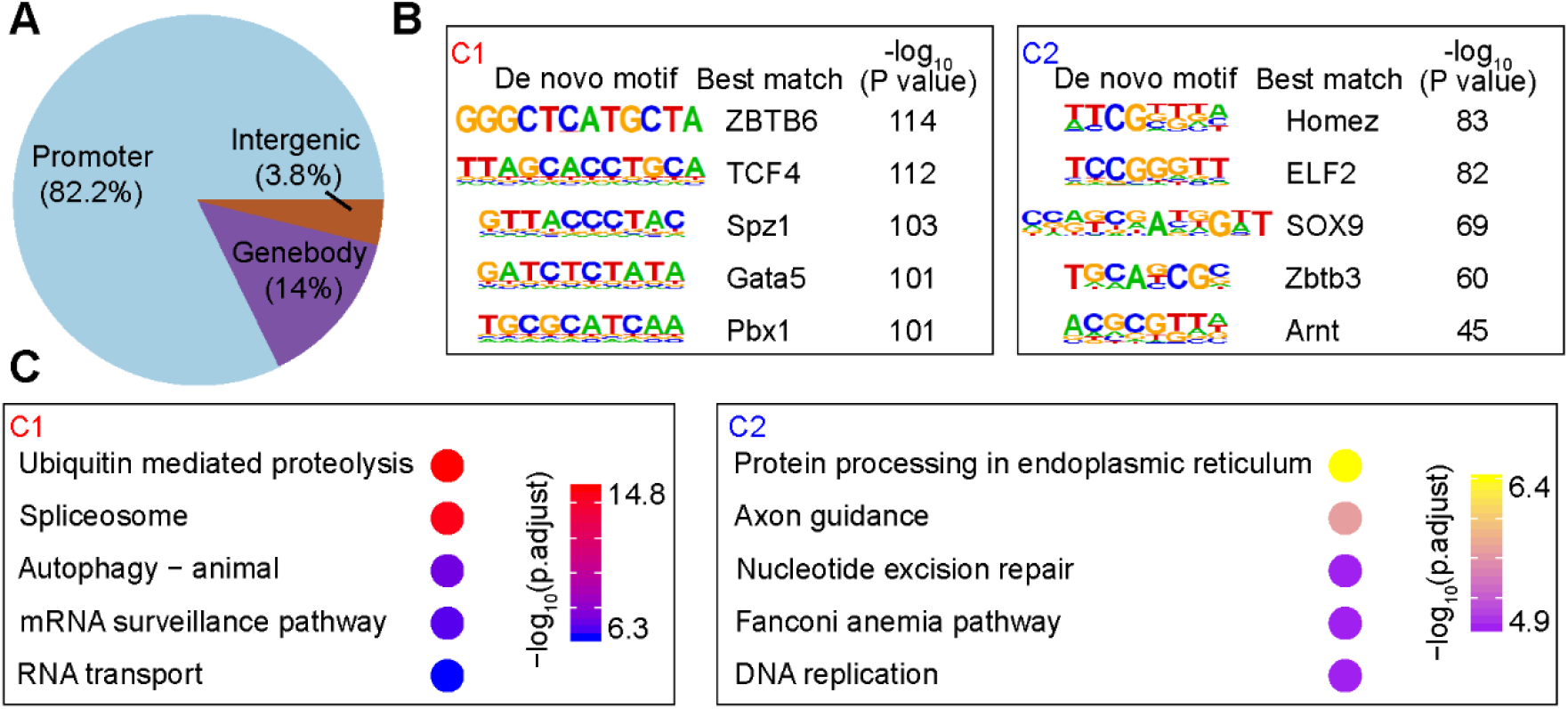
Functional analysis of H3K9cr peaks in the embryonic forebrain. (A) Pie chart showing distribution of H3K9cr peaks at annotated genomic regions. (B) Sequence logos corresponding to enriched elements identified by de novo motif analysis of two groups of H3K9cr peaks at promoter regions. (C) KEGG pathway enrichment analysis of genes an-notated in these peaks.

### Genome-wide alterations of histone Kcr under metabolic stimulation

To profile dynamic changes of histone Kcr at the genome level, we used anti-pan-Kcr antibody to capture Kcr signal of core histone [7, 26]. Kcr could be detected in Nestin^+^ NSPCs in ventricular zone (VZ) and subventricular zone (SVZ) of the developing mouse brains (S3A Fig). We next performed anti-pan-Kcr ChIP assay in cultured NSPCs, and found that histone Kcr was also positively correlated with gene expression in NSPCs (S3B amd S3C Fig). *In vitro*, treatment with crotonate (a crotonyl-CoA donor for crotonylation) or MS-275 (a selective inhibitor of class I HDACs) resulted in elevated histone Kcr level (Fig 3A and 3B), while C646 (a selective inhibitor of p300) significantly inhibited the increase of histone Kcr (Fig 3C). We also observed an increased histone Kcr level in the embryonic forebrain at E13.5 under treatment of MS-275 (S3D amd S3E Fig).

**Fig 3.**
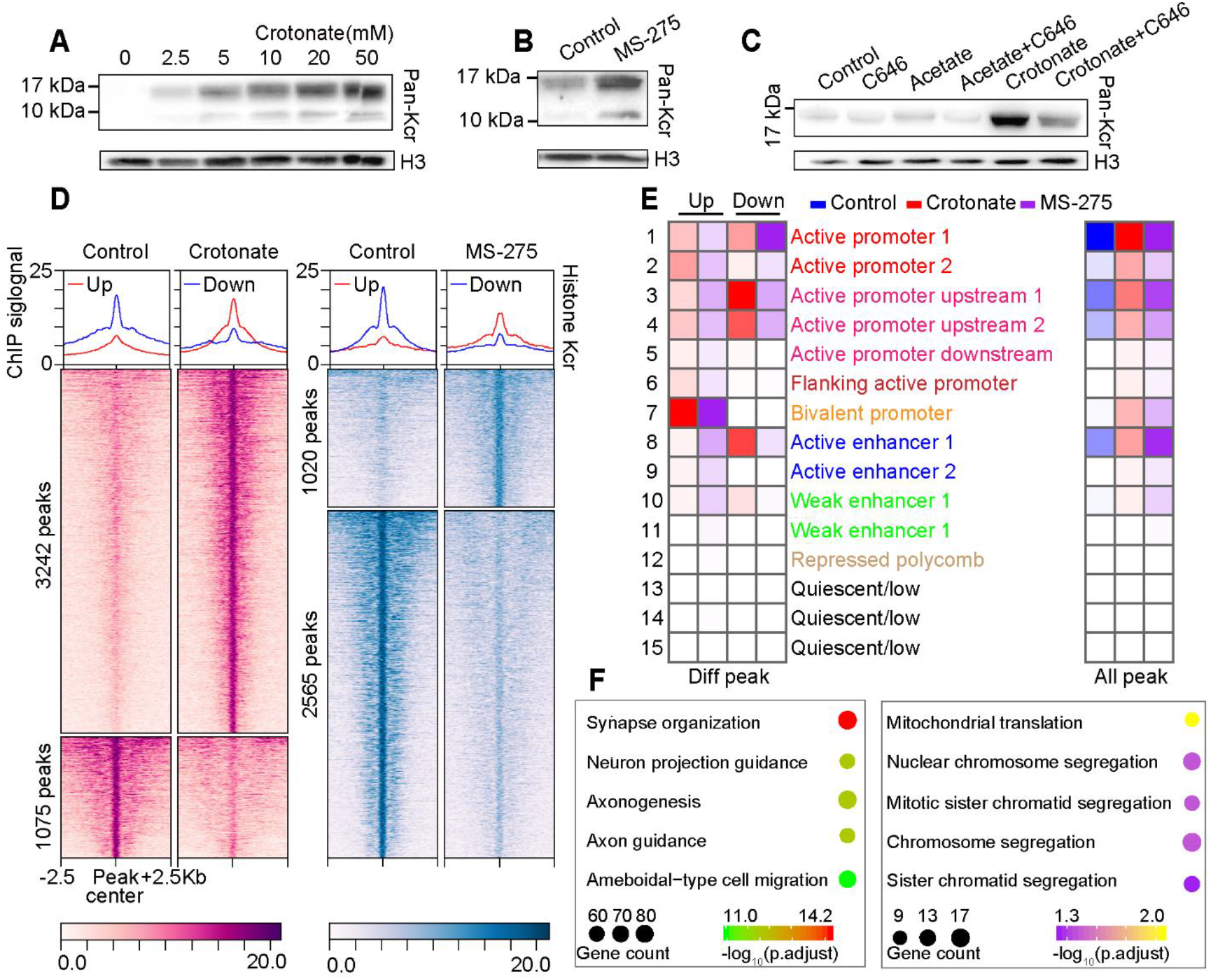
Dynamic changes of histone Kcr in NSPCs. (A) Crotonate resulted in substantially elevated histone Kcr level in cultured NSPCs. (B) Western blotting analysis of histone Kcr level in cultured NSPCs after indicated treatment. (C) MS-275 affects core histone modifications in NSPCs. (D) Average profiles of histone Kcr peaks divided into increased enrichment (Up) and decreased enrichment (Down) group after crotonate or MS-275 treatment. (E) Epigenomic distribution of different groups of peaks. (F) GO analysis of genes annotated in these peaks with increased enrichment (left panel) and decreased enrichment (right panel) after crotonate treatment.

We then conducted a quantitative comparison of ChIP-seq data (crotonate vs control, and MS-275 vs control) to assess dynamic changes of histone Kcr in NSPCs. We divided different binding peaks into two groups, the “Up” group consisting of peaks with increased histone Kcr enrichment, and the “Down” group containing peaks with decreased histone Kcr enrichment after crotonate or MS-275 treatment (Fig 3D and S3F Fig). Interestingly, most peaks in the “Up” group were located at bivalent promoters, the crucial cis-regulatory elements involving in transcriptional regulation of development related genes [27]. while peaks of the “Down” group were enriched in proximal active promoter regions (Fig 3E). Gene ontology (GO) enrichment analysis of genes annotated in these peaks indicated an over-representation of genes involved in cell-fate determination of NSPCs, suggesting a potential role of histone Kcr in the metabolic regulation of neural compartment (Fig 3F). Together, we propose that histone Kcr may remodel local chromatin environment through its dynamic changes at promoter or enhancer regions under metabolic stimulation.

### Histone Kcr activates bivalent promoters to stimulate gene expression

Although histone Kcr has been previously shown to facilitate transcription in a cell-free system [8], genome-wide direct association of histone Kcr with gene expression remains largely unknown. To address this question, we took three independent approaches to perform integrated analysis of ChIP-seq and RNA-seq data of NSPCs. Firstly, binding and expression target analysis (BETA) showed that histone Kcr peaks with increased enrichment had significant effects as a gene activator rather than repressor under crotonate treatment (Fig 4A) [28]. Secondly, we performed fisher’s exact tests comparing genes that contain differential histone Kcr enrichment at promoters to those that are differentially expressed. We found significant correlations between histone Kcr enrichment and altered gene expression, particularly for genes that display increased expression (Fig 4B; and S4A Fig). We finally performed gene set enrichment analysis (GSEA) using gene sets that display different histone Kcr binding at promoters. We found that crotonate-induced changes in histone Kcr level were significantly associated with concordant changes in gene expression (Fig 4C, S4B and S4C Fig).

**Fig 4.**
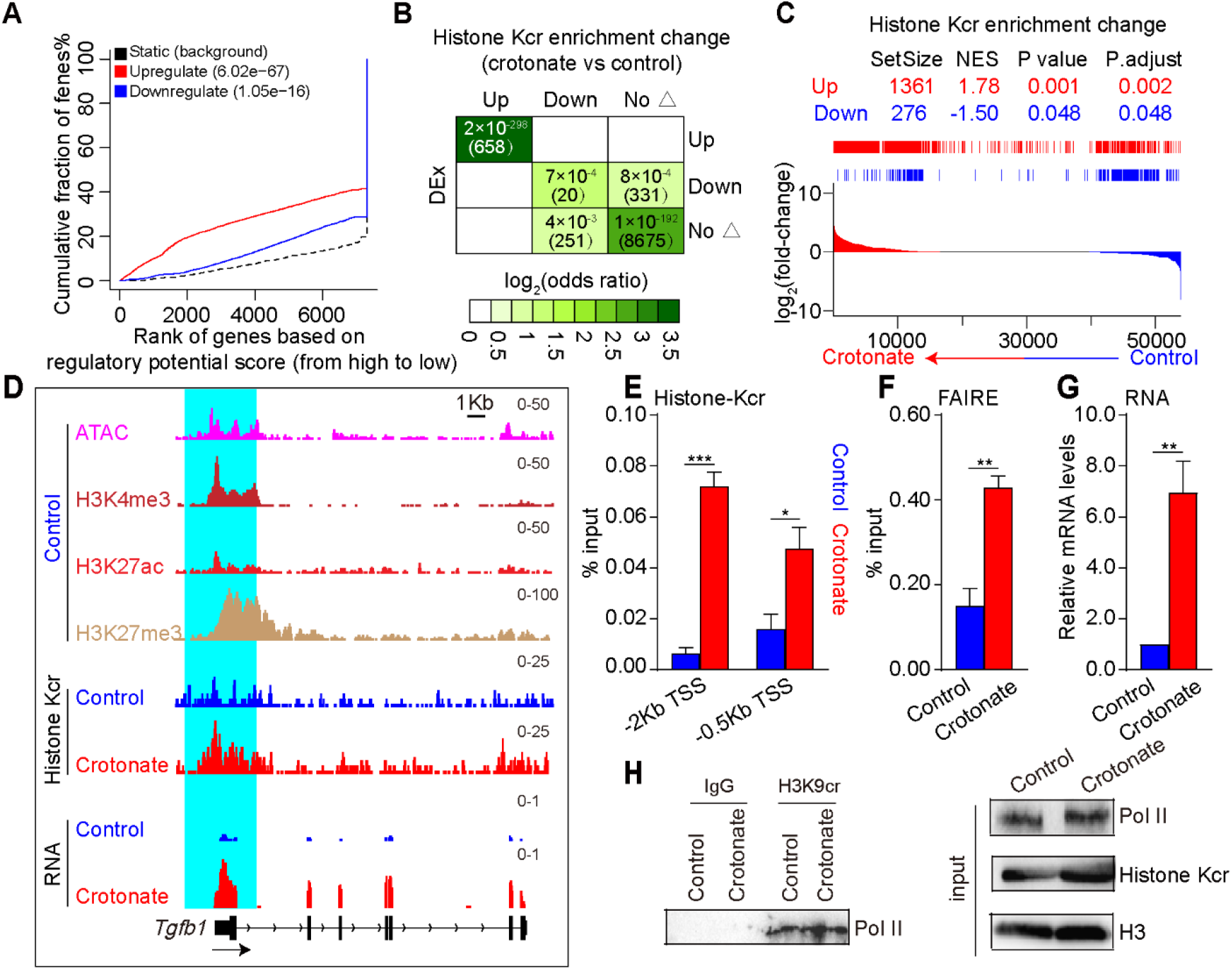
Histone Kcr activates bivalent promoters to promote gene expression. (A) BETA plot of combined computational analysis of histone Kcr ChIP-seq and RNA-seq data (peaks with increased histone Kcr enrichment as input, crotonate vs control). (B) Odds-ratio analysis of overlapping genes displaying differential histone Kcr enrichment versus differential gene expression (DEx), insert numbers indicate respective P-values for associations, with the number of genes overlapping in parentheses. (C) GSEA analysis of association between changes of histone Kcr at promoters and gene expression changes. (D) Genome-browser view at *Tgfb1* gene of different sequencing data sets. Proximal promoter region (+/-2Kb of TSS) was highlighted in a cyan background. (E) ChIP-qPCR result of histone Kcr at the *Tgfb1* promoter. (F) FAIRE-qPCR result of chromatin openness at the *Tgfb1* promoter. (G) qRT-PCR result of gene expression of *Tgfb1*. (H) Co-IP showing Pol II binding at H3K9cr enriched regions. Unpaired t-test was conducted to analyze statistical significance in E-G. Error bars represent SE, ***P< 0.001.

*Tgfb1* and *Notum* are genes under control of bivalent promoters in NSPCs, and the levels of both histone Kcr enrichment at their promoters and expression of these genes were increased significantly after crotonate treatment (Fig 4D and S4D Fig). Therefore, we selected these two genes to validate sequencing data and elucidate the mechanisms underlying histone Kcr-mediated activation of bivalent promoters. Indeed, our ChIP-qPCR and formaldehyde-assisted isolation of regulatory elements followed by qPCR analysis (FAIRE-qPCR) data indicated that crotonate treatment significantly increased histone Kcr enrichment and chromatin openness at both promoters (Fig 4E, 4F, S4E and S4F Fig), accompanied by a significant upregulation of gene expression (Fig 4G and S4G Fig). In contrast, minor changes in histone Kac were observed after crotonate treatment compared with MS-275 treatment (S4H Fig). Moreover, crotonate predominantly resulted in an increase of histone Kcr versus histone Kac (S4I Fig), even though there was a comparable ability of histone Kcr to stimulate gene expression compared with histone Kac (S4J and S4K Fig). It is worth noting that Pol II binding was also observed at regions with H3K9cr binding (Fig 4H), suggesting potentially coordinated recruitment of transcription machinery with histone Kcr. Overall, these results provide strong evidence suggesting that histone Kcr not only serves as an epigenetic mark that is highly correlated with gene expression, but also as a potentially mechanistic way to coordinate transcriptional activation via modulating chromatin openness and recruitment of transcriptional machinery.

### Histone Kcr regulates NSPCs fate via remodeling transcriptome

To further explore the cellular functions of histone Kcr, we examined the effects of crotonate on neuronal cell proliferation. By using bromodeoxyuridine (BrdU) pulse-labelling, we found that less BrdU was incorporated in NSPCs treated with crotonate than control treatment (Fig 5A). Importantly, after 3-day differentiation, NSPCs treated with crotonate differentiated into more TUJ1^+^ neurons than the control group (Fig 5B). To link transcriptome changes with the cell-fate determination, we integrated our bulk RNA sequencing data with published single-cell RNA sequencing data from two independent studies on embryonic neurogenesis [29, 30]. A total of 22 cell clusters were identified in E14.5 cortex (S5A and S5B Fig). Specific cell markers were used to distinguish radial glia (RG), progenitors (SVZ), and neurons (excitatory), as well as proliferative cells (P) and non-proliferative cells (Np) (S5C-5E Fig). We found that down-regulated genes upon crotonate treatment were enriched in the category which can switch NSPCs from the state of proliferation to the state of exit from mitosis and thus promote a loss of “stemness” or undifferentiated state (Fig 5C, 5D and S6A-S6H Fig). On the other hand, up-regulated genes were mainly related to the category that promotes differentiation of NSPCs into neurons (Fig 5D, S6G and S6H Fig), and shows a tendency to fall into neuronal-type state (N_type_) (S6I Fig) [30].

**Fig 5.**
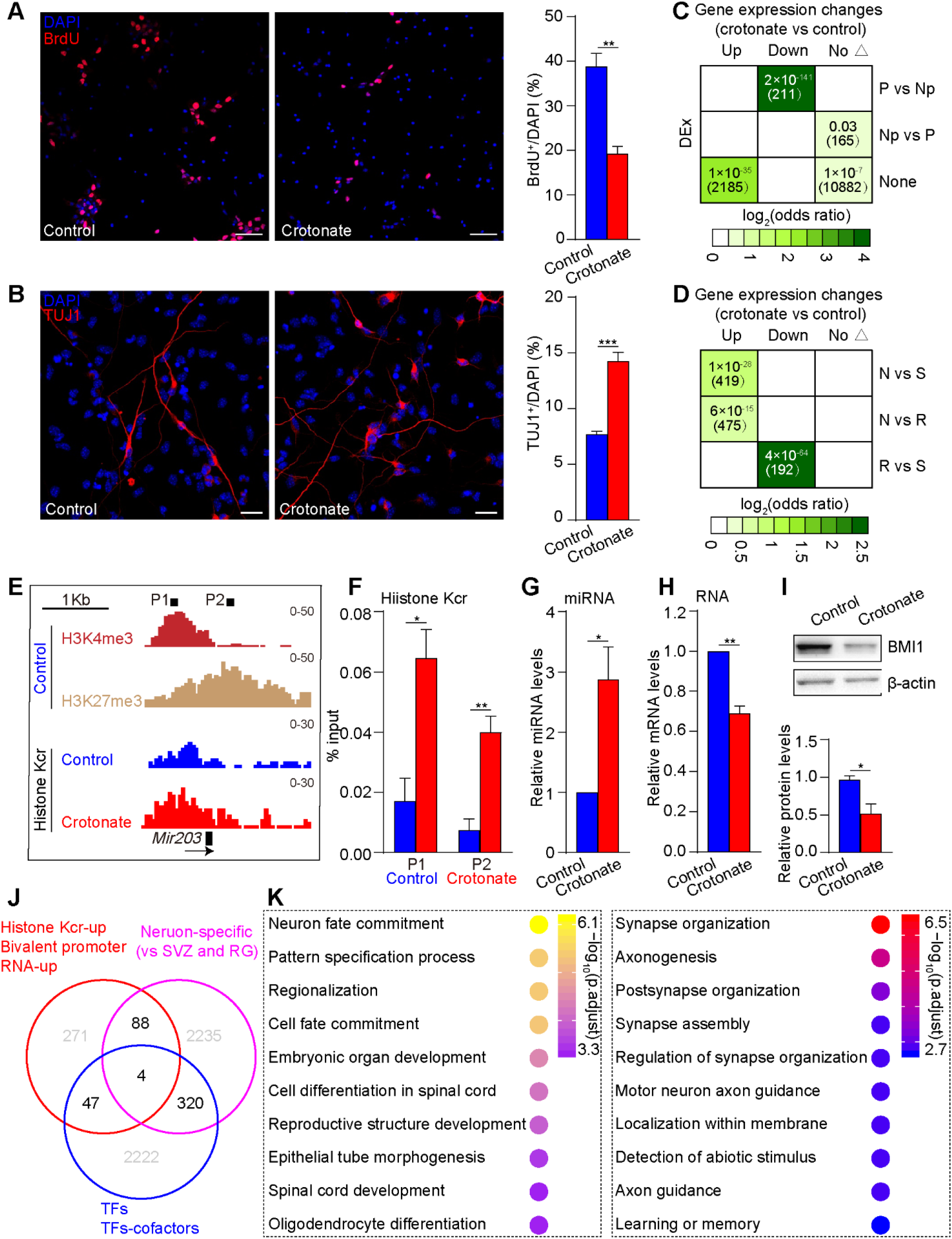
Histone Kcr alters NSPCs fate via transcriptome remodeling. (A) Left panel, representative images illustrating reduced proliferation ability of NSPCs. Scale bar, 50 μm. Right panel, statistical analysis for BrdU incorporation rate. (B) Left panel, representative images illustrating enhanced differentiation ability of NSPCs. Scale bar, 20 μm. Right panel, statistical analysis of TUJ1^+^ of NSPCs derived neurons. (C-D) Odds-ratio analysis of overlapping genes displaying differential gene expression (DEx) versus representing changes of cell fate, insert numbers indicate respective P-values for associations, with the number of genes overlapping in parentheses. (E) Genome-browser view at *Mir-203* gene of different sequencing data sets. (F) ChIP-qPCR result of histone Kcr at the *Mir-203* proximal regions, the exact positions of the 2 primer pairs are depicted in E. (G-(H) qRT-PCR result of gene expression of *Mir-203* and *Bmi1*. (I) Western blotting result of changes in Bmi1 protein levels after cotonate treatment. (J) Venn diagram of a combined comparison of up-regulated genes annotated in bivalent promoters with increased histone Kcr enrichment (red circle, M > 0 with P-values < 0.05), genes representing neuronal differentiation (purplish red circle) and genes encoding mouse transcription factors (TFs) and TFs-cofactors (blue circle). (K) GO analysis of biological processes of those overlapping genes. Left pane, genes intersected in red circle and blue circle; right panel, genes intersected in red circle and purplish red circle. Unpaired t-test was conducted to analyze statistical significance in A-B and F-I. Error bars represent SE, ***P< 0.001.

Interestingly, miR-203, a microRNA we have recently found to inhibit proliferation via *Bmi1* repression in NSPCs [31], was under regulation of bivalent promoters and dis-played an increased histone Kcr level at its proximal promoter regions, as well as upregulation of expression after crotonate treatment in NSPCs (Fig 5E-5G and S7A-C Fig). In consistent with our previous findings, a reduction of both mRNA and protein levels of Bmi1 was observed concordant with up-regulation of miR-203 (Fig 5H and 5I). In contrast, we found sharp declines of histone Kcr at promoters of mitosis-related genes and histone genes that are important for S phase (S7D-S7F Fig). Interestingly, genes that contain activated bivalent promoters upon crotonate treatment are enriched in neuronal fate determination and maturation, such as *Socs2* and *Pacsin1* (Fig 5J, 5K, S8A and S8B Fig). Of note, the effects of crotonate on histone acylation metabolism were not limited to histone Kcr (S8C-S8E Fig). Taken together, our molecular and functional characterization of histone Kcr suggests that activation of bivalent promoters via histone Kcr may be a key regulatory mechanism for cell-fate determination of NSPCs.

## Discussion

Although histone Kcr has been previously shown to localize at promoters and potential enhancers of active genes [7], up to now, detailed patterns of distribution and dynamic changes of histone Kcr at the epigenomic level remain still unknown. Here, to our knowledge, we provide the first investigation on genome-wide distribution and dynamic changes of histone Kcr and its correlation with gene expression. We explored the potential role of histone Kcr as a key regulatory mechanism underlying gene activation and disturbed NSPCs fate. Under normal conditions, histone Kcr marks active promoters with high level of H3K4me3, H3K27ac and open chromatin configuration. These highly expressed genes are involved in nucleic acid and protein metabolism as well as cell proliferation. However, under crotonate-stimulating conditions, crotonate-derived crotonyl-CoA may break the balance of crotonylation and decrotonylation at local chromatin environment which may in turn facilitate p300/CBP-mediated deposition of histone Kcr at bivalent promoters and thus activate expression of genes otherwise silenced in NSPCs. This is likely through enhancing chromatin openness and binding of Pol II. Contributing to evident transcriptome remodeling, these activated genes promote NSPCs transition from stemness state to a state that tends to favor neuronal differentiation (Fig 6).

**Fig 6.**
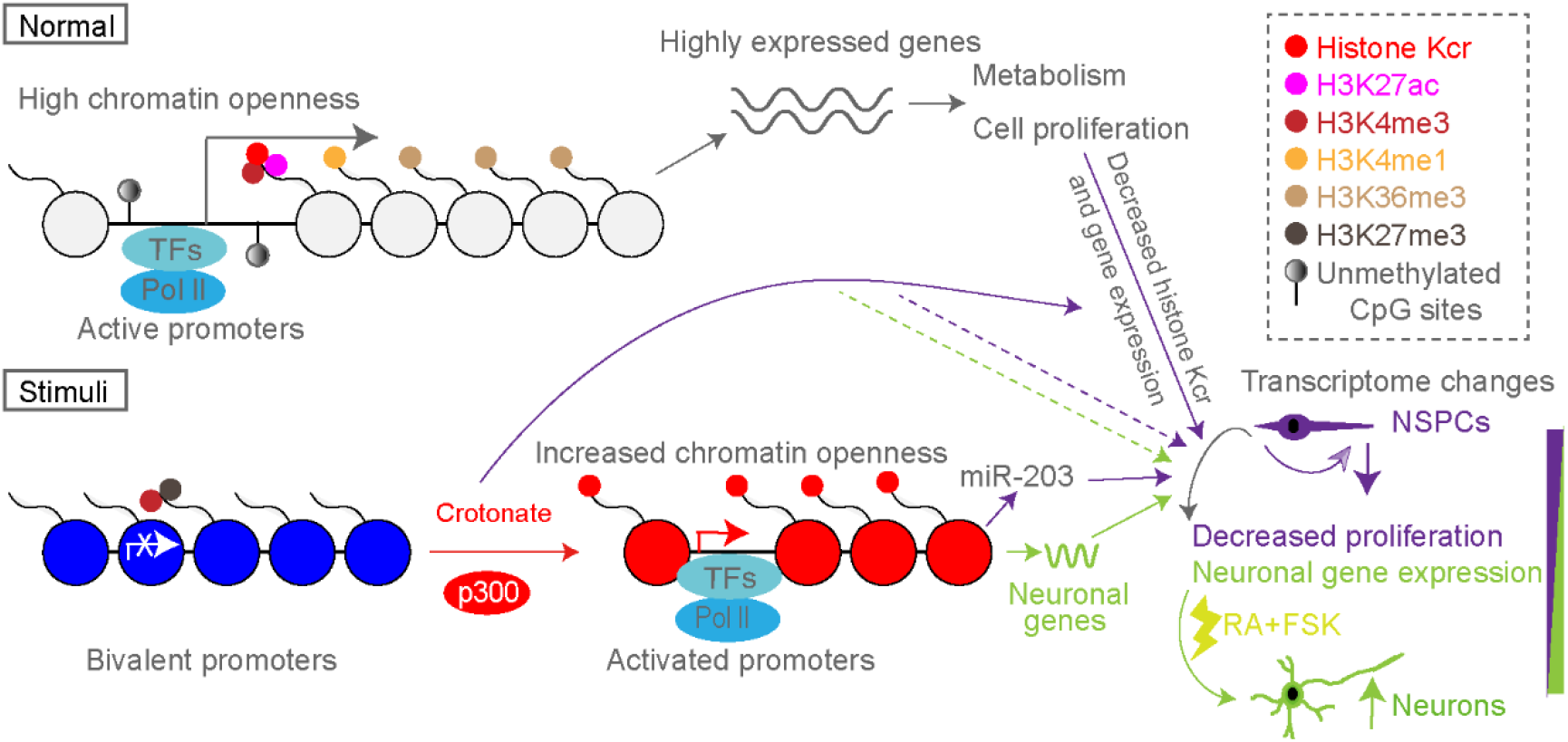
A proposed model of histone Kcr in regulating proliferation and differentiation of NSPCs.

It remains to be defined how histone Kcr regulates transcription. In previous studies, histone Kcr has been considered as a stimulator for transcription mainly through preferential binding enhancement of histone Kcr over Kac, under certain circumstances such as HDAC1-VRPP overexpression, CDYL deletion and AF9 (F59A) mutation [8, 10, 11, 17]. However, two recent studies have revealed roles of histone Kcr in transcriptional repression. Notably, in response to an increase of histone Kcr, down-regulated genes displayed a simultaneous decrease of histone Kac in the metabolic cycle of yeast and NEAT-knockdown cells [32, 33]. Therefore, the balance between the opposite changes of histone Kcr and Kac, accompanied by the complex crosstalk among histone PTMs, makes it difficult and ambiguous to determine whether histone Kcr directly represses transcription. In general, increased histone Kcr enrichment leads to a decrease in H3K27me3 and an enhanced recognition by reader modules, like AF9, which bridge the transcriptional machinery to local loci and gene transcription [13, 17]. Here we provide evidence supporting an important role of histone Kcr for enhancing chromatin openness and binding of Pol II in NSPCs. These findings lead us to propose that histone Kcr may play a causative, not just correlative, role for activation of bivalent promoters and transcriptional regulation, which is consistent with the chemical properties of histone Kcr such as altering strong electrostatic interactions between oppositely charged nucleosome DNA [34].

Metabolism plays important roles in cell-fate determination in part through epigenetic regulation, especially during neurogenesis [35]. Acetate, a precursor of acetyl-CoA, delays differentiation of ESCs and blocks early histone deacetylation in a dose-dependent manner [36]. Crotonate is converted to crotonyl-CoA through ACSS2, which acts as a donor for crotonylation on histones and non-histone proteins, and also stimulates histone Kcr through recruitment of histone crotonyltransferase [8, 37, 38]. To our knowledge, we report, for the first time, the role of histone Kcr in NSPCs at the metabolic-epigenetic level. Using published single-cell sequencing data of embryonic neurogenesis, we show that crotonate-treated NSCPs differentiate into neurons at a higher ratio (Fig 5B). Our mechanistic studies suggest that crotonate-induced increase in histone Kcr promotes transcriptome remodeling which favors the switch from stemness state to neuron-like state via activation of bivalent promoters. Although crotonate has been reported to inhibit proliferation of cancer cells and regulate histone Kcr levels in cell cycle, the mechanism by which histone Kcr controls cell proliferation remains unclear [19, 37]. We propose that the histone Kcr-miR-203-Bmi1 regulatory axis may play a critical role in regulating proliferation of NSPCs. Importantly, the sharp decline of histone Kcr at the promoter of histone genes as well as genes involved in progression of M phase, provides an alternative role for histone Kcr to regulate proliferation. In this scenario, a decrease of histone Kcr may set a “brake” of gene expression that is important for cell cycle progression in response to signals that disfavor proliferation. However, our results do not exclude the possibility that crotonate may induce other effects apart from histone Kcr to contribute to transcriptome changes and cell-fate determination of NSPCs. In addition, our current studies only analysed histone Kcr at promoter regions and it is possible that binding of histone Kcr at enhancers and other regulatory regions may also be involved in the regulation of gene expression.

Overall, we provide the first genome-wide analysis of histone Kcr in neural cells and identify a unique role of histone Kcr in gene activation and cell fate determination. Our findings may promote a deeper understanding of histone Kcr in the context of epigenetic regulation during neural development and diseases, as well as offer a new perspective on activation of local chromatin caused by these metabolically sensitive histone lysine acylations.

## Materials and methods

### Ethical statement

All mice used in the current study are in a C57BL6 background. All mouse experiments were approved by the Animal Committee of the Institute of Zoology, Chinese Academy of Sciences, Beijing, China. Mice were housed in groups of 3–5 animals in a 12h:12h light: dark cycle, with standard mouse chow and water ad libitum.

### Cell culture

NSPCs isolated from the embryonic forebrain at E13.5 were cultured in N2 medium [DMEM/F12 medium (Invitrogen) supplemented with 20 ng/ml epidermal growth factor and 20 ng/ml fibroblast growth factor (EGF/FGF, PeproTech), 1% penicillin/streptomycin (Invitrogen), 0.5× B27 (Invitrogen), and 0.5× N2 (Invitrogen)]. Neurons isolated from P0 mouse hippocampus were cultured in Neurobasal medium (Invitrogen) supplemented with 1% penicillin/streptomycin, 1% B27, and 2 mM GlutaMax (Invitrogen) at 5% CO2, 37°C incubator as previously described [39]. For passaging, bulk NSPCs were dissociated mechanically to single cells and were then sub-cultured.

### Analysis of NSPCs proliferation and differentiation

To study cell proliferation, NSPCs were dissociated using mechanical pipettes, and then plated on poly-L-ornithin/laminin coated glass coverslips at a density of 3×10^4^ cells/well in the N2 medium. BrdU pulse-labelling was used to assess proliferation of NSPCs. For differentiation analyses, NSPCs were seeded at a density of 5×10^4^ cells/well, and forskolin (FSK, Sigma-Aldrich) and retinoic acid (RA, Sigma-Aldrich) were employed to induce differentiation of NSPCs.

### Immunocytochemistry (ICC) and immunostaining

Immunocytochemistry and Immunostaining was performed as previously described [31]. DAPI (Sigma-Aldrich) was used to label the cell nucleus. The primary and secondary antibodies used were listed in Table S1 of S1 Data. All images were acquired by confocal microscopy (ZEISS, LSM710).

### Western blotting and co-immunoprecipitation

Proteins were separated using sodium dodecyl sulfate-polyacrylamide gel electrophoresis (SDS-PAGE) and transferred to polyvinylidene fluoride membranes (Millipore). After incubated with antibodies, the immunoreactive products were detected by using Tanon-5200 Chemiluminescent Imaging System (Tanon, China, Shanghai). Co-immunoprecipitation was performed according to the instructions of the Protein A/G PLUS-Agarose (Santa Cruz).

### ChIP-seq, ChIP-qPCR and FAIRE-qPCR

ChIP was performed as described previously [40]. Briefly, the nuclear lysates of NSPCs which were fixed in 1% formaldehyde were sonicated to an average size approximately 200–600 bp. Normal rabbit IgG, Pan-Kcr and Pan-Kac antibodies were used for immunoprecipitation reaction. ChIP samples were sequenced on BGISEQ-500 and subjected to qRT-PCR on Roche LightCycler^®^ 480 II. The FAIRE-qPCR procedure was adapted from a previously published protocol [41].

### RNA-seq and qRT-PCR

RNA was extracted using TRIzol^®^ reagent according to the instructions (Invitrogen). RNA samples were sequenced on Illumina NovaSeq 6000 platform and subjected to qRT-PCR on Roche LightCycler^®^ 480 II.

### ChIP-seq, RNA-seq and Single-cell RNA-seq data analysis

ChIP-seq files were analyzed using the following tools: Bowtie2 for read mapping [42], MACS2 for peak calling [43], and MAnorm for differential binding analyze [44]. RNA-seq FASTQ files were analyzed using the following tools: Salmon (v.1.0.0) for read mapping [45], and DEseq2 for differential gene expression analysis [46]. Seurat was implemented to analyze single-cell RNA-seq data [47]. Detailed analysis steps are provided within Supplementary methods of S1 Data.

## Supporting information

**S1 Data. PDF containing Supplementary tables and methods.**

**S1 Fig.**
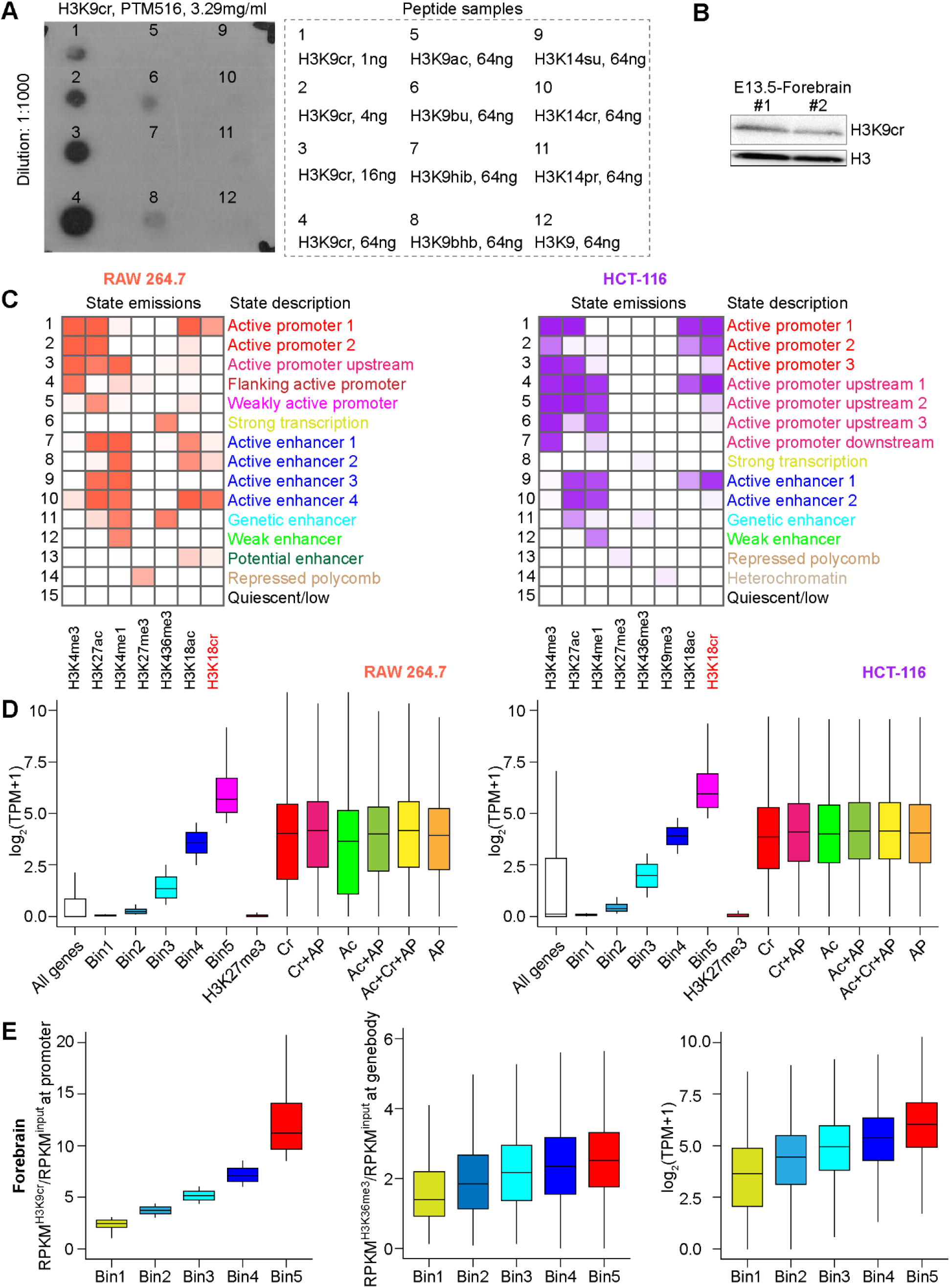
Genome-wide profiling and analysis of histone Kcr. (A) Dot plot assay of H3K9cr antibody demonstrating its specificity, with peptide samples on the right panel. (B) Western blotting analysis of H3K9cr expression in the embryonic (E13.5) forebrain. (C) Heatmap showing distribution of H3K18cr signal on fifteen chromatin states modelled using ChromHMM in RAW 264.7 (left panel) and HCT-116 (right panel). (D) Boxplots showing gene expression changes of different groups of genes: TPM were divided into five bins, as indicated, from lowly expressed genes (Bin1) to highly expressed genes (Bin5); genes under control of indicated histone marks (Cr: H3K18cr; Ac: H3K18ac; AP: H3K4me3+H3K27ac, active promoters) in RAW 264.7 (left panel) and HCT-116 (right panel). (E) Left panel, boxplot showing changes of promoter histone Kcr level of different groups of genes, ratio of histone Kcr signal to input signal at promoter regions (+/-1 kb of TSS) was divided into five bins, as indicated, from genes with low ratio (Bin1) to genes with high ratio (Bin5) in forebrain. Middle panel, boxplot showing changes of gene body H3K36me3 level of different groups of genes defined previously. Right panel, boxplot showing gene expression changes of different groups of genes defined previously.

**S2 Fig.**
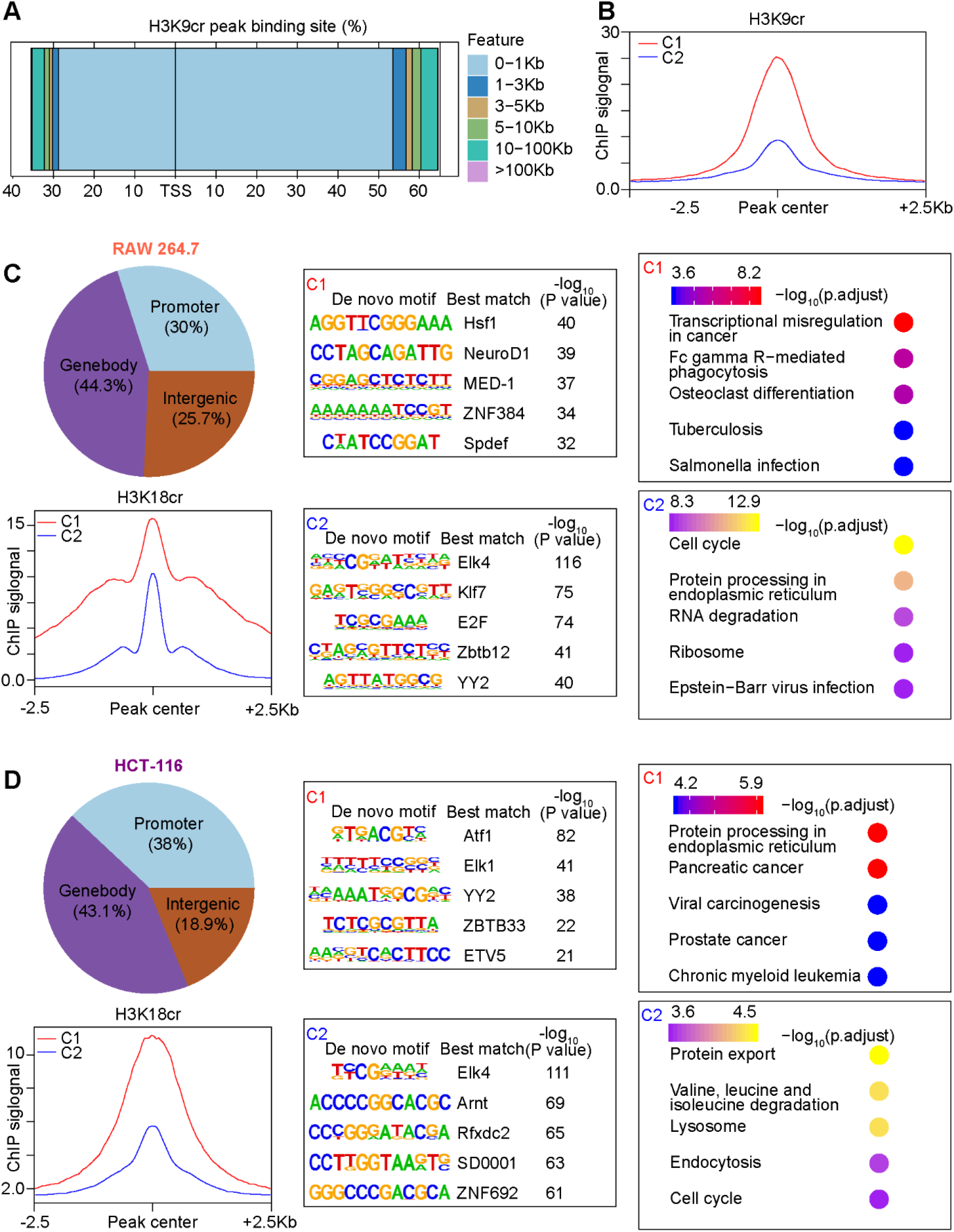
Distribution and functional analysis of histone Kcr peaks. (A) Bar plot showing distribution of H3K9cr peak to TSS. (B) Average profile of H3K9cr within +/-2.5 kb of center of H3K9cr peaks at promoter regions which were divided into two clusters by H3K9cr signal (C1-C2). (C-D) Pie chart showing distribution of H3K18cr peaks, and average profiles, motif analyze of those H3K18cr peaks at promoter regions, as well as KEGG pathway enrichment analysis of genes annotated at promoter peaks in RAW 264.7 (C) and HCT-116 (D).

**S3 Fig.**
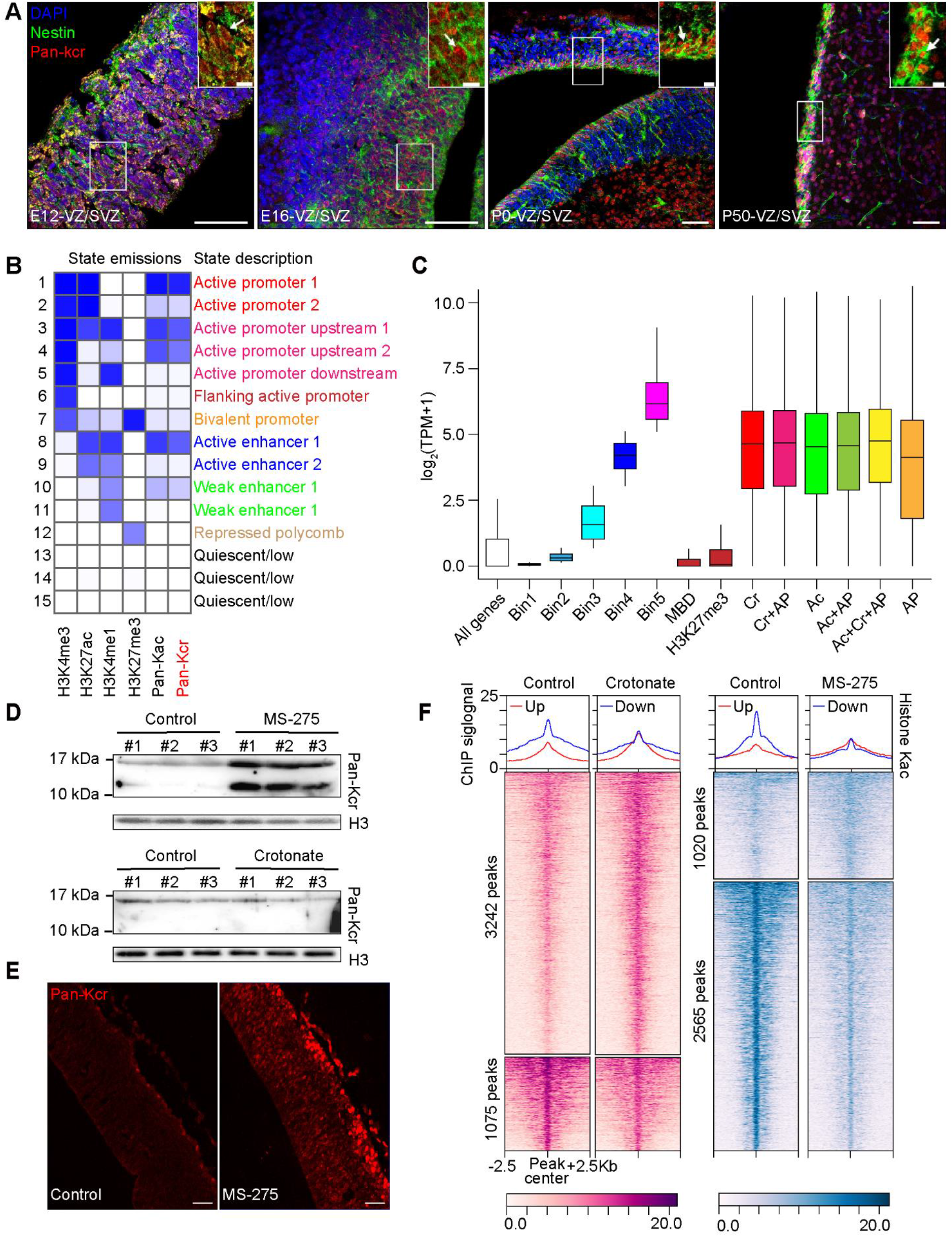
Histone Kcr distribution and changes in NSPCs. (A) Fluorescent staining of Kcr on mouse forebrain sections at E12, E16, P0, and P50 revealed its high expression in Nestin^+^ NSPCs in the VZ/SVZ. Magnified images of boxed areas are shown in the upperright corners. White arrowheads indicate Kcr^+^ Nestin^+^ NSPCs. Scale bar, 50 μm; Scale bar in magnified box, 10 μm. (B) Heatmap showing distribution of histone Kcr signal on fifteen chromatin states modelled using ChromHMM in NSPCs. (C) Boxplot showing gene expression changes of different groups of genes: TPM of genes were divided into five bins, as indicated, from lowly expressed genes (Bin1) to highly expressed genes (Bin5); genes under control of indicated histone marks (Cr: Histone Kcr; Ac: Histone Kac; AP: H3K4me3+H3K27ac, active promoters) in NSPCs. (D) MS-275 affects histone Kcr level of the embryonic forebrain in vivo. Top panel, western blotting analysis indicated that histone Kcr level was affected by MS-275 abundance. In the experiment, 0.1% DMSO was intraperitoneally injected as a control, and 20 mg/kg MS-275 was injected every 24 h for continuous three days on pregnant mice at Day 9.5 of pregnancy. Bottom panel, western blotting analysis indicated crotonate could not induce changes of histone Kcr level in vivo. PBS was intraperitoneally injected as a control, and 500 μM / 30 g crotonate was injected every 24 h for continuous three days on pregnant mice at Day 9.5 of pregnancy. (E) Immunostaining results showed MS-275 induced histone Kcr changes in VZ/SVZ at E13.5 stage. Scale bar, 50 μm. (F) Average profiles of histone Kac over peaks of “Up” group and “Down” group defined in Fig 3D.

**S4 Fig.**
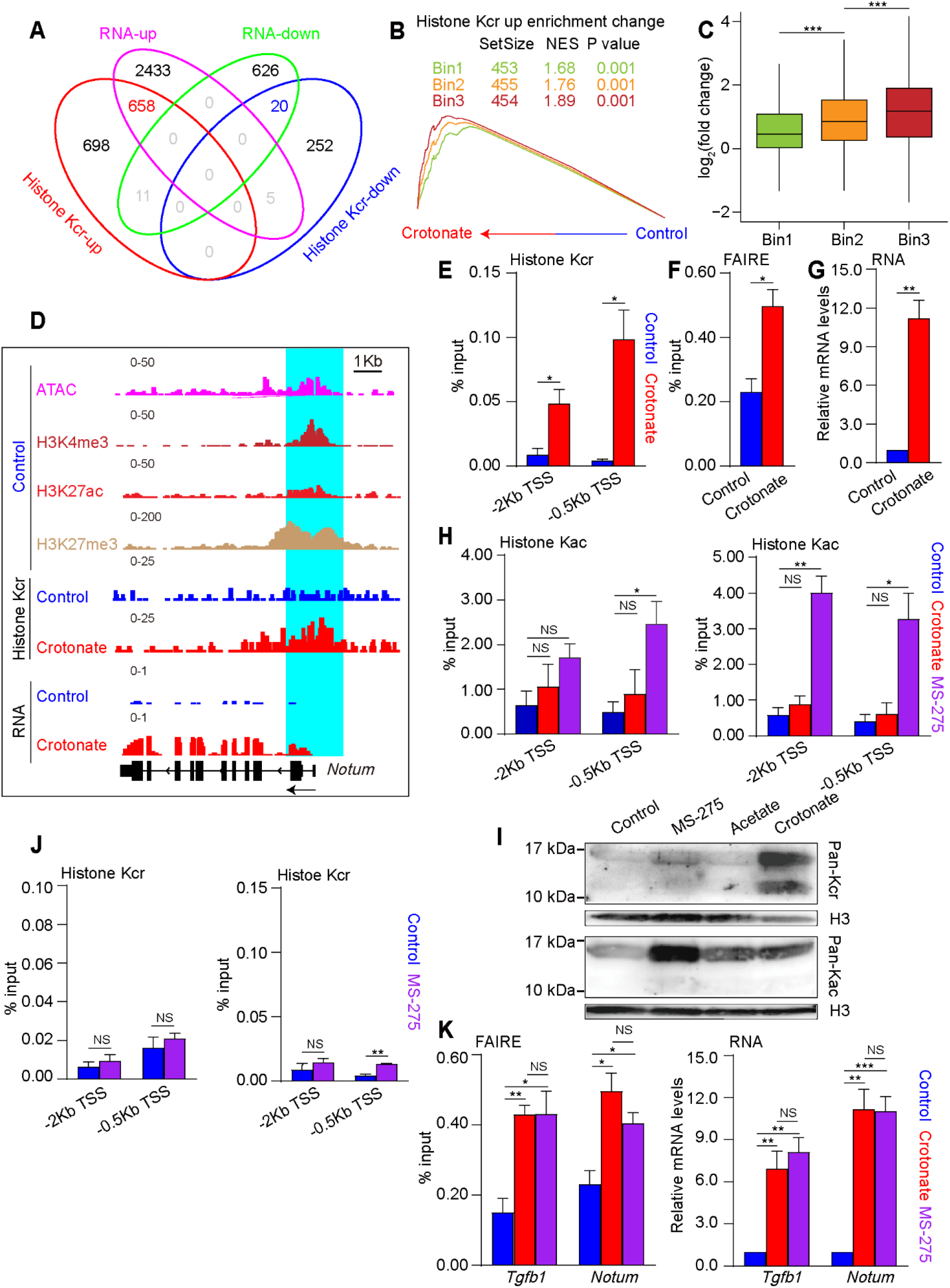
Crotonate stimulates gene expression by histone Kcr. (A) Venn diagram of a combined comparison of differentially expressed genes (RNA-up: up-regulated genes; RNA-down: down-regulated genes) and genes with different promoter histone Kcr binding (Histone Kcr-up: genes with increased histone Kcr enrichment; Histone Kcr-down: genes with decreased histone Kcr enrichment). (B) GSEA analysis of association between changes of histone Kcr enrichment at promoter and gene expression changes. Genes with increased promoter histone Kcr enrichment were divided into three bins, as indicated, with extent of increase form low (Bin1) to high (Bin3). (C) Boxplot showing gene expression changes of different groups of genes defined in B, unpaired wilcox test was conducted to calculate the statistical significances, ***P< 0.001. (D) Genome-browser view at *Notum* gene of different sequencing data sets, proximal promoter region (+/-2Kb of TSS) was highlighted in a cyan background. (E) ChIP-qPCR result of histone Kcr at the *Notum* promoter. (F) FAIRE-qPCR result of chromatin openness at the *Notum* promoter. (G) qRT-PCR result of gene expression of *Notum*. (H) ChIP-qPCR result of histone Kac at the *Tgfb1* promoter (left panel) and the *Notum* promoter (right panel). (I) Western blotting analysis of histone Kcr level in cultured NSPCs under indicated treatment. (J) ChIP-qPCR result of histone Kcr at the *Tgfb1* promoter (left panel) and the *Notum* promoter (right panel). (K) FAIRE-qPCR result of chromatin openness at the *Tgfb1* promoter and the *Notum* promoter (left panel), and qRT-PCR result of gene expression of *Tgfb1* and *Notum* (right panel). Unpaired t-test was conducted to analyse statistical significance in E-H and J, K Error bars represent SE, ***P< 0.001.

**S5 Fig.**
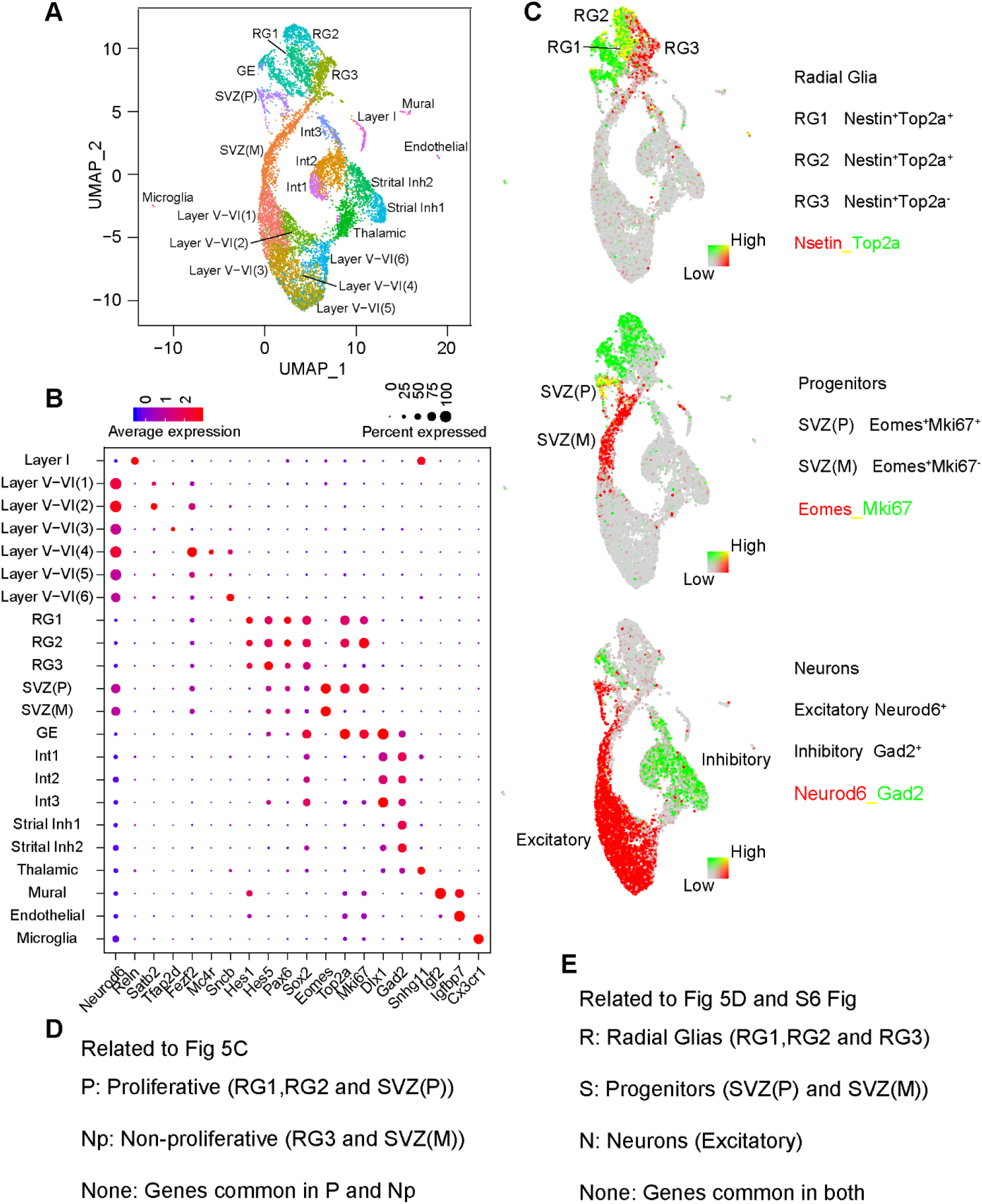
Defining various cell types in embryonic cortex using published single-cell RNA sequencing data. (A) Uniform manifold approximation and projection (UMAP) visualization of the 22 clusters identified in E14.5 cortex. (B) Dot plot showing representative markers used to distinguish different cell types in A. (C) Visualization of E14.5 cells and overlaid with gene expression information of canonical marker genes, Nestin (radial glia), Tbr2 (progenitors), Neurond6 (excitatory neurons), Gad2 (inhibitory neurons), and Top2a/Mki67 (proliferating cells). The expression is depicted from gray (low) to red or green (high). (D-E) Abbreviation and corresponding content used in Fig 5 and S6 Fig

**S6 Fig.**
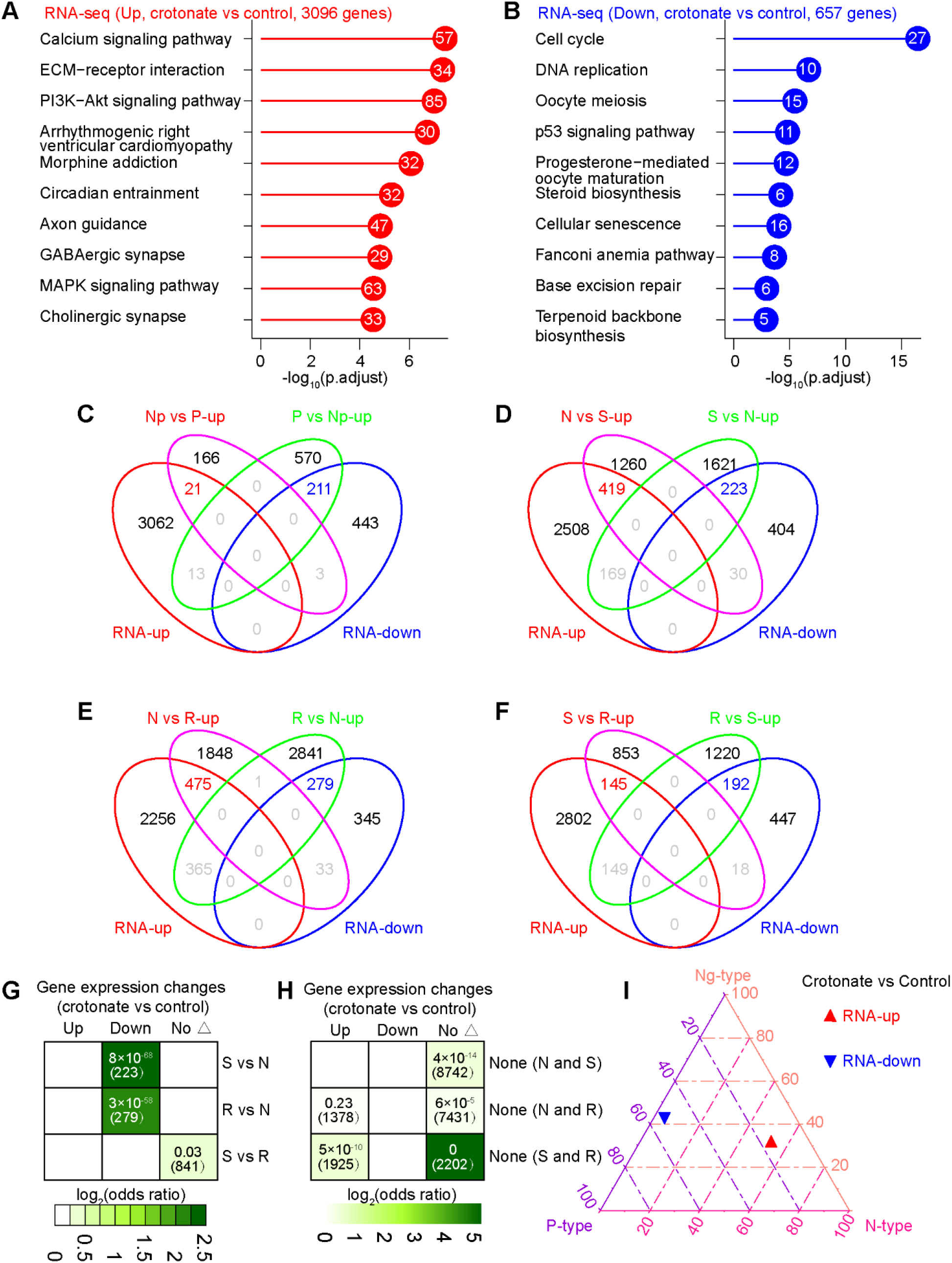
Transcriptome remodeling in crotonate treated NSPCs. (A-B) KEGG pathway enrichment analysis of up-regulated genes (A) and down-regulated genes (B) after crotonate treatment in NSPCs. (C-F) Venn diagram of a combined comparison of genes displaying differential gene expression and representing changes of cell fate. (G, H) Odds-ratio analysis of overlapping genes displaying differential gene expression (DEx) versus representing changes of cell fate, insert numbers indicate respective P-values for associations, with the number of genes overlapping in parentheses. (I) Ternary plot showing transcriptome distributions of up-regulated genes (RNA-up) and down-regulated genes (RNA-down) on different cell state in neurogenesis after crotonate treatment in NSPCs.

**S7 Fig.**
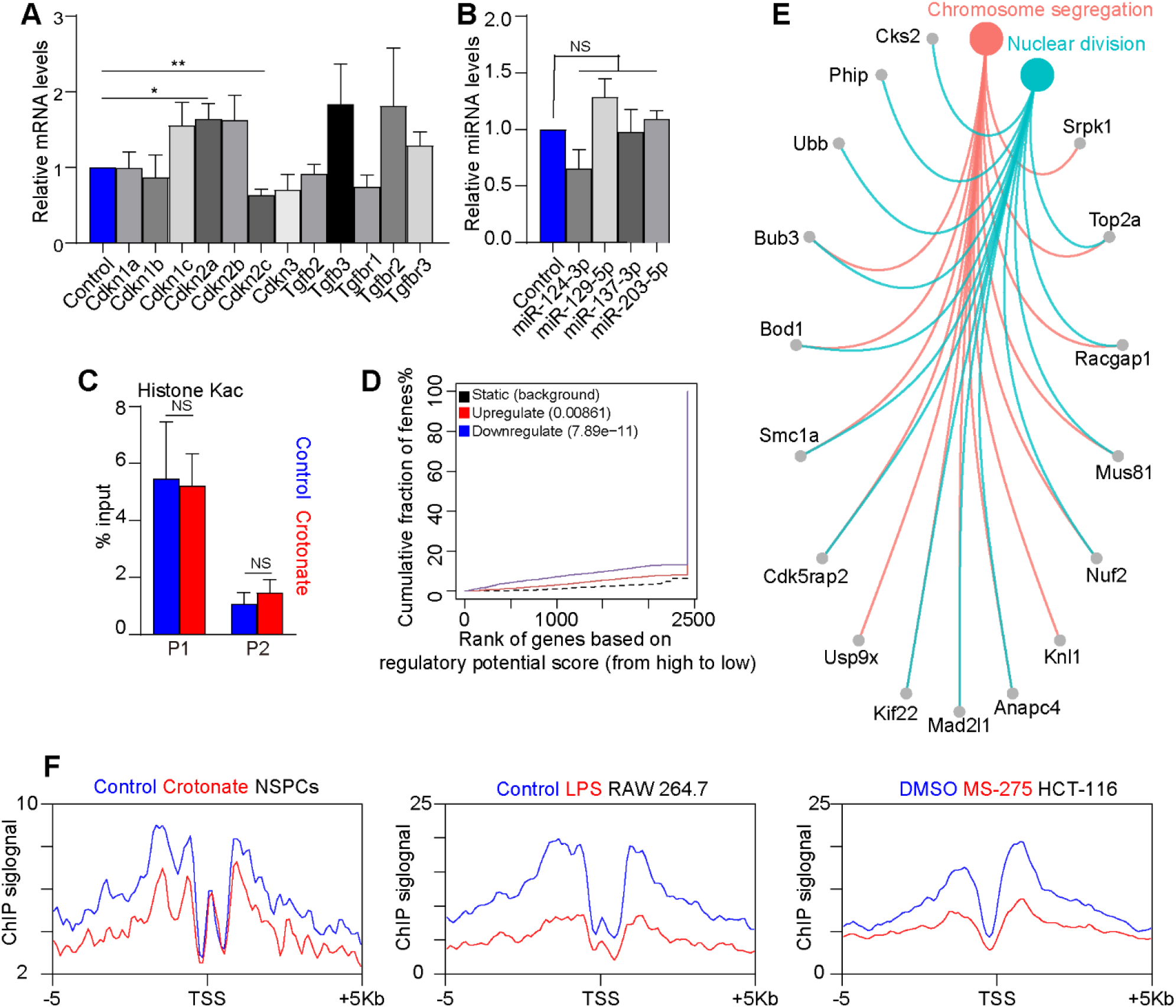
Association of histone Kcr with cell proliferation. (A) qRT-PCR result of gene expression after crotonate treatment. (B) qRT-PCR result of different miRNAs expression which were reported to involve in regulating NSPCs proliferation. (C) ChIP-qPCR result of histone Kac at the *Mir-203* proximal regions, the exact positions of the 2 primer pairs are depicted in Fig 5E. (D) BETA plot of combined computational analysis of histone Kcr ChIP-seq and RNA-seq data (peaks with decreased histone Kcr enrichment as input, crotonate vs control). (E) GO enrichment analysis of genes with decreased histone Kcr enrichment at promoters and gene expression (134 genes, M-values < 0 with P-values < 0.05). (F) Average profiles of histone Kcr within +/-5 kb of TSS of *Hist/HIST* gene cluster.

**S8 Fig.**
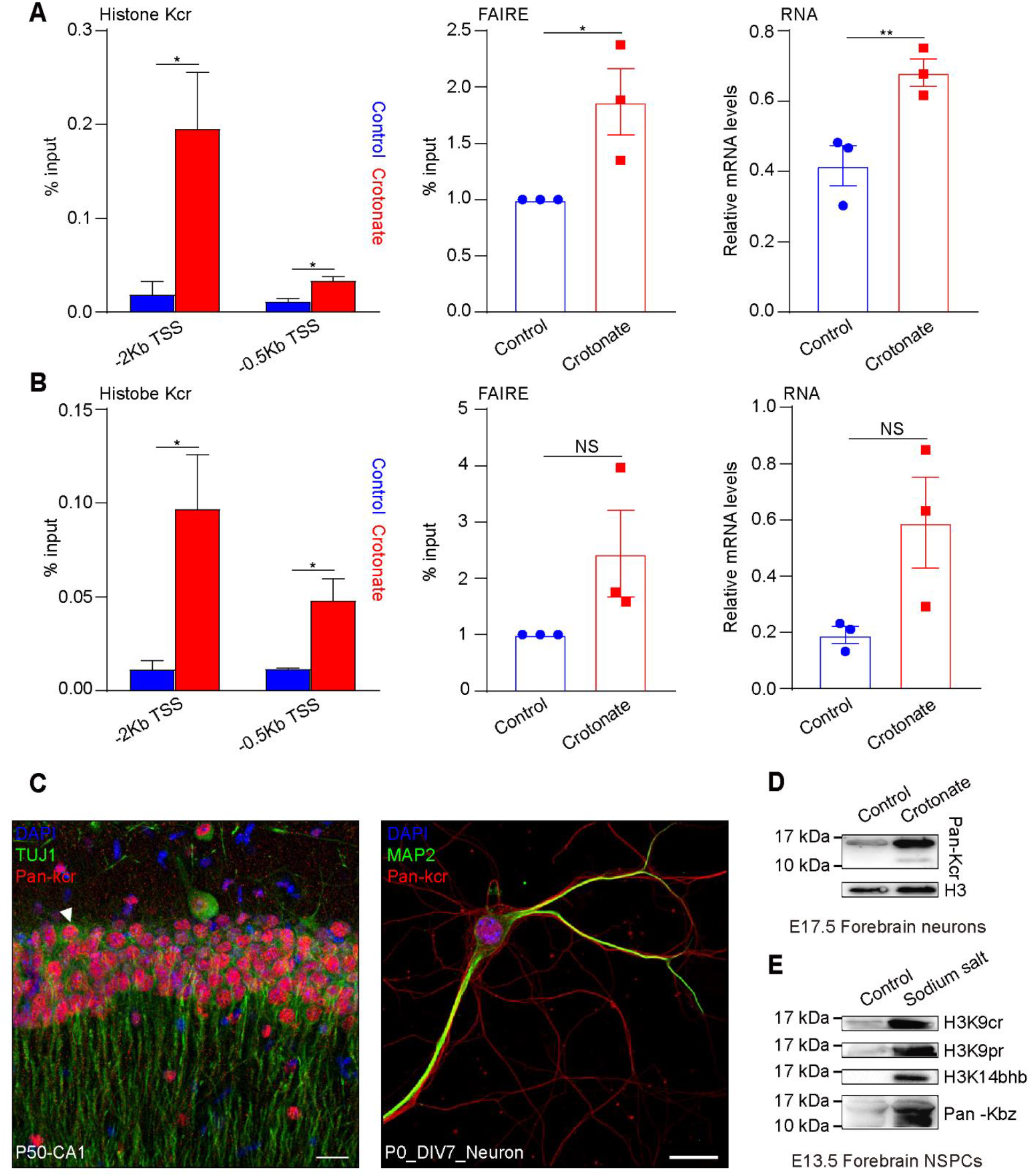
Association of histone Kcr with neuronal differentiation. (A, B) Left panel, ChIP-qPCR result of histone Kcr at the *Socs2* (A) *and Pacsin1* (B) promoter. Middle panel, FAIRE-qPCR result of chromatin openness at the *Socs2* (A) *and Pacsin1* (B) promoter. Right panel, qRT-PCR result of gene expression of *Socs*2 (A) and Pacsin1 (B). (C) Left panel, image of co-localization of Kcr with neuronal marker TUJ1 in vivo, Scale bar, 20 μm. Right panel, immunocytochemical staining of Kcr and neuronal dendritic marker MAP2 in vitro. Scale bar, 50 μm. (D) Crotonate resulted in increase in histone Kcr level after 10 mM crotonate treatment for continuous three days in cultured neurons. (E) Western blotting analysis of histone acylations level in NSPCs which were treated with 10 mM crotonate (H3K9cr), sodium propionate (H3K9pr), sodium 3-hydroxybutanoate (H3K14bhb) and sodium benzoate (Pan-Kbz) separately.

## Acknowledgements

This work was supported by the Strategic Priority Research Program of the Chinese Academy of Sciences (XDA16010300), the National Key Research and Development Program of China Project (2016YFA0101402) and grants from the National Science Foundation of China (91753140, 81771224).

## References

1. Li X, Egervari G, Wang Y, Berger SL, Lu Z. Regulation of chromatin and gene expression by metabolic enzymes and metabolites. Nat Rev Mol Cell Biol. 2018;19(9):563–78. Epub 2018/06/23. doi: 10.1038/s41580-018-0029-7. PubMed PMID: 29930302.

2. Zentner GE, Henikoff S. Regulation of nucleosome dynamics by histone modifications. Nat Struct Mol Biol. 2013;20(3):259–66. Epub 2013/03/07. doi: 10.1038/nsmb.2470. PubMed PMID: 23463310.

3. Lawrence M, Daujat S, Schneider R. Lateral Thinking: How Histone Modifications Regulate Gene Expression. Trends Genet. 2016;32(1):42–56. doi: 10.1016/j.tig.2015.10.007. PubMed PMID: WOS:000368214800004.

4. Barnes CE, English DM, Cowley SM. Acetylation & Co: an expanding repertoire of histone acylations regulates chromatin and transcription. Essays Biochem. 2019;63(1):97–107. Epub 2019/04/04. doi: 10.1042/EBC20180061. PubMed PMID: 30940741; PubMed Central PMCID: PMCPMC6484784.

5. Sabari B, Zhang D, Allis C, Zhao Y. Metabolic regulation of gene expression through histone acylations. Nat Rev Mol Cell Biol. 2017;18(2):90–101.

6. Tan M, Luo H, Lee S, Jin F, Yang JS, Montellier E, et al. Identification of 67 histone marks and histone lysine crotonylation as a new type of histone modification. Cell. 2011;146(6):1016–28. Epub 2011/09/20. doi: 10.1016/j.cell.2011.08.008. PubMed PMID: 21925322; PubMed Central PMCID: PMCPMC3176443.

7. Tan M, Luo H, Lee S, Jin F, Yang J, Montellier E, et al. Identification of 67 histone marks and histone lysine crotonylation as a new type of histone modification. Cell. 2011;146(6):1016–28. doi: 10.1016/j.cell.2011.08.008. PubMed PMID: WOS:000295258100022.

8. Sabari BR, Tang ZY, Huang H, Yong-Gonzalez V, Molina H, Kong HE, et al. Intracellular Crotonyl-CoA Stimulates Transcription through p300-Catalyzed Histone Crotonylation. Mol Cell. 2015;58(2):203–15. doi: 10.1016/j.molcel.2015.02.029. PubMed PMID: WOS:000353222900004.

9. Ruiz-Andres O, Sanchez-Nino MD, Cannata-Ortiz P, Ruiz-Ortega M, Egido J, Ortiz A, et al. Histone lysine crotonylation during acute kidney injury in mice. Dis Model Mech. 2016;9(6):633–45. Epub 2016/04/30. doi: 10.1242/dmm.024455. PubMed PMID: 27125278; PubMed Central PMCID: PMCPMC4920150.

10. Wei W, Liu X, Chen J, Gao S, Lu L, Zhang H, et al. Class I histone deacetylases are major histone decrotonylases: evidence for critical and broad function of histone crotonylation in transcription. Cell Res. 2017;27(7):898–915. Epub 2017/05/13. doi: 10.1038/cr.2017.68. PubMed PMID: 28497810; PubMed Central PMCID: PMCPMC5518989.

11. Liu SM, Yu HJ, Liu YQ, Liu XH, Zhang Y, Bu C, et al. Chromodomain Protein CDYL Acts as a Crotonyl-CoA Hydratase to Regulate Histone Crotonylation and Spermatogenesis. Mol Cell. 2017;67(5):853–+. doi: 10.1016/j.molcel.2017.07.011. PubMed PMID: WOS:000411128900013.

12. Jiang GC, Nguyen D, Archin NM, Yukl SA, Mendez-Lagares G, Tang YY, et al. HIV latency is reversed by ACSS2-driven histone crotonylation. J Clin Invest. 2018;128(3):1190–8. doi: 10.1172/Jci98071. PubMed PMID: WOS:000431900100028.

13. Liu Y, Li M, Fan M, Song Y, Yu H, Zhi X, et al. Chromodomain Y-like protein–mediated histone crotonylation regulates stress-induced depressive behaviors. Biol Psychiatry. 2019;85(8):635–49.

14. Liu XG, Wei W, Liu YT, Yang XL, Wu J, Zhang Y, et al. MOF as an evolutionarily conserved histone crotonyltransferase and transcriptional activation by histone acetyltransferase-deficient and crotonyltransferase-competent CBP/p300. Cell Discov. 2017;3:17016. doi: UNSP 17016 10.1038/celldisc.2017.16. PubMed PMID: WOS:000414915100001.

15. Andrews FH, Shinsky SA, Shanle EK, Bridgers JB, Gest A, Tsun IK, et al. The Taf14 YEATS domain is a reader of histone crotonylation. Nat Chem Biol. 2016;12(6):396–U33. doi: 10.1038/nchembio.2065. PubMed PMID: WOS:000376160600007.

16. Zhao D, Guan HP, Zhao S, Mi WY, Wen H, Li YY, et al. YEATS2 is a selective histone crotonylation reader. Cell Research. 2016;26(5):629–32. doi: DOI 10.1038/cr.2016.49. PubMed PMID: WOS:000377449200011.

17. Li Y, Sabari BR, Panchenko T, Wen H, Zhao D, Guan H, et al. Molecular Coupling of Histone Crotonylation and Active Transcription by AF9 YEATS Domain. Mol Cell. 2016;62(2):181–93. Epub 2016/04/23. doi: 10.1016/j.molcel.2016.03.028. PubMed PMID: 27105114; PubMed Central PMCID: PMCPMC4841940.

18. Xiong X, Panchenko T, Yang S, Zhao S, Yan P, Zhang W, et al. Selective recognition of histone crotonylation by double PHD fingers of MOZ and DPF2. Nat Chem Biol. 2016;12(12):1111–8. Epub 2016/10/25. doi: 10.1038/nchembio.2218. PubMed PMID: 27775714; PubMed Central PMCID: PMCPMC5253430.

19. Fellows R, Denizot J, Stellato C, Cuomo A, Jain P, Stoyanova E, et al. Microbiota derived short chain fatty acids promote histone crotonylation in the colon through histone deacetylases. Nature Communications. 2018;9(1):105. doi: ARTN 105 10.1038/s41467-017-02651-5. PubMed PMID: WOS:000419658100005.

20. Kelly RDW, Chandru A, Watson PJ, Song Y, Blades M, Robertson NS, et al. Histone deacetylase (HDAC) 1 and 2 complexes regulate both histone acetylation and crotonylation in vivo. Sci Rep-Uk. 2018;8(1):14690. doi: ARTN 14690 10.1038/s41598-018-32927-9. PubMed PMID: WOS:000446035900020.

21. Tang T, Zhang Y, Wang Y, Cai Z, Lu Z, Li L, et al. HDAC1 and HDAC2 Regulate Intermediate Progenitor Positioning to Safeguard Neocortical Development. Neuron. 2019;101(6):1117–33 e5. Epub 2019/02/03. doi: 10.1016/j.neuron.2019.01.007. PubMed PMID: 30709655.

22. Falkenberg KJ, Johnstone RWJNrDd. Histone deacetylases and their inhibitors in cancer, neurological diseases and immune disorders. Nat Rev Drug Discov. 2014;13(9):673–91.

23. Hwang AW, Trzeciakiewicz H, Friedmann D, Yuan CX, Marmorstein R, Lee VMY, et al. Conserved Lysine Acetylation within the Microtubule-Binding Domain Regulates MAP2/Tau Family Members. PLoS One. 2016;11(12):e0168913. doi: ARTN e0168913 10.1371/journal.pone.0168913. PubMed PMID: WOS:000392853100076.

24. Scott CE, Wynn SL, Sesay A, Cruz C, Cheung M, Gomez Gaviro MV, et al. SOX9 induces and maintains neural stem cells. Nat Neurosci. 2010;13(10):1181–9. Epub 2010/09/28. doi: 10.1038/nn.2646. PubMed PMID: 20871603.

25. Forrest MP, Hill MJ, Quantock AJ, Martin-Rendon E, Blake DJ. The emerging roles of TCF4 in disease and development. Trends Mol Med. 2014;20(6):322–31. Epub 2014/03/07. doi: 10.1016/j.molmed.2014.01.010. PubMed PMID: 24594265.

26. Lu Y, Xu Q, Liu Y, Yu Y, Cheng ZY, Zhao Y, et al. Dynamics and functional interplay of histone lysine butyrylation, crotonylation, and acetylation in rice under starvation and submergence. Genome Biol. 2018;19(1):144. Epub 2018/09/27. doi: 10.1186/s13059-018-1533-y. PubMed PMID: 30253806; PubMed Central PMCID: PMCPMC6154804.

27. Voigt P, Tee WW, Reinberg D. A double take on bivalent promoters. Genes Dev. 2013;27(12):1318–38. Epub 2013/06/22. doi: 10.1101/gad.219626.113. PubMed PMID: 23788621; PubMed Central PMCID: PMCPMC3701188.

28. Wang S, Sun H, Ma J, Zang C, Wang C, Wang J, et al. Target analysis by integration of transcriptome and ChIP-seq data with BETA. Nat Protoc. 2013;8(12):2502–15. Epub 2013/11/23. doi: 10.1038/nprot.2013.150. PubMed PMID: 24263090; PubMed Central PMCID: PMCPMC4135175.

29. Loo L, Simon JM, Xing L, McCoy ES, Niehaus JK, Guo J, et al. Single-cell transcriptomic analysis of mouse neocortical development. Nat Commun. 2019;10(1):134. Epub 2019/01/13. doi: 10.1038/s41467-018-08079-9. PubMed PMID: 30635555; PubMed Central PMCID: PMCPMC6329831.

30. Telley L, Govindan S, Prados J, Stevant I, Nef S, Dermitzakis E, et al. Sequential transcriptional waves direct the differentiation of newborn neurons in the mouse neocortex. Science. 2016;351(6280):1443–6. Epub 2016/03/05. doi: 10.1126/science.aad8361. PubMed PMID: 26940868.

31. Liu PP, Tang GB, Xu YJ, Zeng YQ, Zhang SF, Du HZ, et al. MiR-203 Interplays with Polycomb Repressive Complexes to Regulate the Proliferation of Neural Stem/Progenitor Cells. Stem Cell Reports. 2017;9(1):190–202. Epub 2017/06/13. doi: 10.1016/j.stemcr.2017.05.007. PubMed PMID: 28602614; PubMed Central PMCID: PMCPMC5511050.

32. Wang Z, Zhao Y, Xu N, Zhang S, Wang S, Mao Y, et al. NEAT1 regulates neuroglial cell mediating Aβ clearance via the epigenetic regulation of endocytosis-related genes expression. Cell Mol Life Sci. 2019:1–14.

33. Gowans GJ, Bridgers JB, Zhang J, Dronamraju R, Burnetti A, King DA, et al. Recognition of Histone Crotonylation by Taf14 Links Metabolic State to Gene Expression. Mol Cell. 2019;76(6):909–21 e3. Epub 2019/11/05. doi: 10.1016/j.molcel.2019.09.029. PubMed PMID: 31676231; PubMed Central PMCID: PMCPMC6931132.

34. Onufriev AV, Schiessel H. The nucleosome: from structure to function through physics. Curr Opin Struc Biol. 2019;56:119–30. doi: 10.1016/j.sbi.2018.11.003. PubMed PMID: WOS:000482514600016.

35. Xie K, Sheppard AJB. Dietary Micronutrients Promote Neuronal Differentiation by Modulating the Mitochondrial-Nuclear Dialogue. Bioessays. 2018;40(7):1800051.

36. Moussaieff A, Rouleau M, Kitsberg D, Cohen M, Levy G, Barasch D, et al. Glycolysis-mediated changes in acetyl-CoA and histone acetylation control the early differentiation of embryonic stem cells. Cell Metab. 2015;21(3):392–402. Epub 2015/03/05. doi: 10.1016/j.cmet.2015.02.002. PubMed PMID: 25738455.

37. Wei W, Mao A, Tang B, Zeng Q, Gao S, Liu X, et al. Large-Scale Identification of Protein Crotonylation Reveals Its Role in Multiple Cellular Functions. J Proteome Res. 2017;16(4):1743–52. Epub 2017/02/25. doi: 10.1021/acs.jproteome.7b00012. PubMed PMID: 28234478.

38. Kollenstart L, de Groot AJ, Janssen GM, Cheng X, Vreeken K, Martino F, et al. Gcn5 and Esa1 function as histone crotonyltransferases to regulate crotonylation-dependent transcription. J Biol Chem. 2019:jbc. RA119. 010302.

39. Tang GB, Zeng YQ, Liu PP, Mi TW, Zhang SF, Dai SK, et al. The Histone H3K27 Demethylase UTX Regulates Synaptic Plasticity and Cognitive Behaviors in Mice. Front Mol Neurosci. 2017;10:267. doi: ARTN 267 10.3389/fnmol.2017.00267. PubMed PMID: WOS:000408345400001.

40. Liu C, Teng ZQ, Santistevan NJ, Szulwach KE, Guo W, Jin P, et al. Epigenetic regulation of miR-184 by MBD1 governs neural stem cell proliferation and differentiation. Cell Stem Cell. 2010;6(5):433–44. Epub 2010/05/11. doi: 10.1016/j.stem.2010.02.017. PubMed PMID: 20452318; PubMed Central PMCID: PMCPMC2867837.

41. Li X, Zhao Q, Wei W, Lin Q, Magnan C, Emami MR, et al. The DNA modification N6-methyl-2’-deoxyadenosine (m6dA) drives activity-induced gene expression and is required for fear extinction. Nat Neurosci. 2019;22(4):534.

42. Langmead B, Salzberg SL. Fast gapped-read alignment with Bowtie 2. Nat Methods. 2012;9(4):357–9. Epub 2012/03/06. doi: 10.1038/nmeth.1923. PubMed PMID: 22388286; PubMed Central PMCID: PMCPMC3322381.

43. Feng JX, Liu T, Qin B, Zhang Y, Liu XS. Identifying ChIP-seq enrichment using MACS. Nature Protocols. 2012;7(9):1728–40. doi: 10.1038/nprot.2012.101. PubMed PMID: WOS:000308526300013.

44. Shao Z, Zhang Y, Yuan GC, Orkin SH, Waxman DJ. MAnorm: a robust model for quantitative comparison of ChIP-Seq data sets. Genome Biol. 2012;13(3):R16. Epub 2012/03/20. doi: 10.1186/gb-2012-13-3-r16. PubMed PMID: 22424423; PubMed Central PMCID: PMCPMC3439967.

45. Patro R, Duggal G, Love MI, Irizarry RA, Kingsford C. Salmon provides fast and bias-aware quantification of transcript expression. Nat Methods. 2017;14(4):417–9. Epub 2017/03/07. doi: 10.1038/nmeth.4197. PubMed PMID: 28263959; PubMed Central PMCID: PMCPMC5600148.

46. Love MI, Huber W, Anders S. Moderated estimation of fold change and dispersion for RNA-seq data with DESeq2. Genome Biol. 2014;15(12):550. Epub 2014/12/18. doi: 10.1186/s13059-014-0550-8. PubMed PMID: 25516281; PubMed Central PMCID: PMCPMC4302049.

47. Stuart T, Butler A, Hoffman P, Hafemeister C, Papalexi E, Mauck WM, 3rd, et al. Comprehensive Integration of Single-Cell Data. Cell. 2019;177(7):1888–902 e21. Epub 2019/06/11. doi: 10.1016/j.cell.2019.05.031. PubMed PMID: 31178118; PubMed Central PMCID: PMCPMC6687398.

